# Overexpression of a Gene That Modulates Cyclic-di-GMP Enhances Granulation in *Mycobacterium smegmatis*

**DOI:** 10.64898/2026.01.25.701641

**Authors:** Tiffany Lam, Samantha J. Belculfine, Joseph G. Gikonyo, Jeffrey J. Kane, Chul Park, Yasu S. Morita

## Abstract

Granulation is a complex microbial-aggregation process essential for forming aerobic granular sludge (AGS) and other microbial granules used in wastewater treatment. However, the biological mechanisms that drive granule formation remain poorly understood. Cyclic-di-GMP (c-di-GMP) is a well-established second messenger that regulates biofilm formation, suggesting it may be used to enhance microbial granulation. *Mycobacterium smegmatis*, a nonpathogenic model bacterium for *Mycobacterium tuberculosis*, naturally forms granules. Because *M. smegmatis* carries a single c-di-GMP modulating gene, *dcpA*, that encodes an enzyme with both diguanylate cyclase (DGC) and phosphodiesterase (PDE) activities, it offers a unique opportunity to examine the role of c-di-GMP in granulation. Here, we generated and studied two engineered *M. smegmatis* strains overexpressing *dcpA* or *dcpA*ΔEAL, the latter of which is defective in PDE activity. Using these engineered strains, we examined different forms of biofilm growth, cell morphology, plastic surface adhesion, granulation, and settleability. Results of sludge volume index and microscopy indicated that the aggregates of *M. smegmatis* were granules rather than flocs, and the settleability of the granules was particularly robust when the cells were grown in a carbon rich medium known to promote granulation. Engineered strains sustained stable granulation more effectively than the wildtype under low concentration Tween-80 treatment, which was used to induce dispersion. These results suggest that overproduction of DcpA and thus the modulated level of intracellular c-di-GMP enhances granulation and promotes granule persistence in *M. smegmatis*. Our study further demonstrates that *M. smegmatis* is a useful model for elucidating biological mechanisms underlying granulation, which could be leveraged to improve granular technologies for wastewater treatment.

## Introduction

Granulation technology for wastewater treatment has grown in popularity over conventional activated sludge because it offers improved sludge settleability and lower capital and operational costs, including reduced energy demand (Hamza et al., 2022; Sepúlveda-Mardones et al., 2019). Some granules such as oxygenic photogranules can also sequester CO_2_ from wastewater and the atmosphere, offering the potential for net-negative-emission wastewater treatment processes (Park and Takeuchi, 2021). Granular systems rely on functional microbial communities that aggregate, degrade organic and inorganic contaminants, and settle rapidly, allowing clarified effluent to be withdrawn from the reactor (Vydehi et al., 2024).

Although granular technology represents a significant advance over conventional activated sludge, several challenges remain, including long start-up periods for granule formation, difficulty maintaining granule size and stability, and the requirement for specialized sequencing batch reactor configurations (Vydehi et al., 2024). These issues likely stem from the limited understanding of the biological mechanisms underlying granulation. Consequently, despite growing interest, wider full-scale adoption of granular technologies has been constrained.

Cyclic-di-GMP (c-di-GMP) is a bacterial second messenger that stimulates production of extracellular polymeric substances (EPS) and suppresses cell motility, thereby promoting biofilm formation (Chen et al., 2025; Cotter and Stibitz, 2007; Isenberg and Mandel, 2024; Purcell and Tamayo, 2016; Römling et al., 2013). Although elevated intracellular c-di-GMP has been linked to increased EPS and enhanced biofilm development, relatively few studies have examined c-di-GMP specifically as a means to promote granulation, a self-aggregated, non-surface-attached form of biofilm that underpins granular technologies (Guo et al., 2017; Shi et al., 2025; Yang et al., 2014). Moreover, previous studies attempted to induce c-di-GMP to promote aggregation by altering operational conditions (*e*.*g*., feeding regimes); however, these indirect, external approaches did not result in consistent c-di-GMP and EPS responses (Guo et al., 2017; Yang et al., 2014). Complicating mechanistic analysis further, wastewater treatment granules comprise complex, mixed environmental consortia.

*Mycobacterium smegmatis* is an environmental heterotroph and nonpathogenic model bacterium for *Mycobacterium tuberculosis* (Sparks et al., 2023). This bacterium can serve as a model to study the mechanisms of microbial granulation for three reasons. First, *M. smegmatis* granulates, and the granulation is enhanced by changes in the nutrient composition of the medium they grow (DePas et al., 2019). Second, like other bacteria, c-di-GMP enhances biofilm formation in *M. smegmatis* (Gupta et al., 2015; Gupta et al., 2016; Ling et al., 2024). Third, unlike other AGS-resident bacteria, mycobacteria possess only a single c-di-GMP modulating gene, *dcpA* (MSMEG_2196). DcpA is a dual-functional enzyme, carrying both diguanylate cyclase (DGC) and phosphodiesterase (PDE) domains responsible for the synthesis and hydrolysis of c-di-GMP, respectively (Bharati et al., 2012; Gupta et al., 2010).

This study aims to genetically modify *M. smegmatis* to test the proof-of-concept that microbial granulation, particularly granulation of heterotrophic bacteria, can be manipulated through intracellular signaling pathways involving c-di-GMP. Through genetic manipulation, we establish that c-di-GMP regulation drives granule formation, providing key mechanistic insights for microbial granulation in wastewater treatment.

## Results

### Characterization of *dcpA* and *dcpA*ΔEAL Overexpression Strains

We cloned the wildtype *dcpA* gene (MSMEG_2196) into the pMV361 expression vector, in which gene expression is driven by a strong Hsp60 promoter in mycobacteria (Stover et al., 1991). An *in vitro* study demonstrated that a point mutation in the EAL motif eliminates the PDE activity of DcpA while maintaining the DGC activity (Bharati et al., 2018). We therefore created another expression vector, in which the EAL motif of DcpA was deleted (*dcpA*ΔEAL).

We first confirmed the expression of *dcpA* and *dcpA*ΔEAL by western blotting. Both proteins are C-terminally tagged with a hemagglutinin (HA) tag. Cell lysates were separated by SDS-PAGE and proteins were detected by anti-HA antibody (**Fig. 1A**). As expected, no band was visible in the wildtype *M. smegmatis* cell lysate. In the lysates of mutant cells expressing *dcpA* or *dcpA*ΔEAL, a single protein band was detected at the expected molecular weight of 68.5 and 68.1 kDa, respectively. DcpA appeared to be more intense than DcpAΔEAL, suggesting that the disruption of the EAL motif in the PDE domain may affect protein stability.

**Figure 1.**
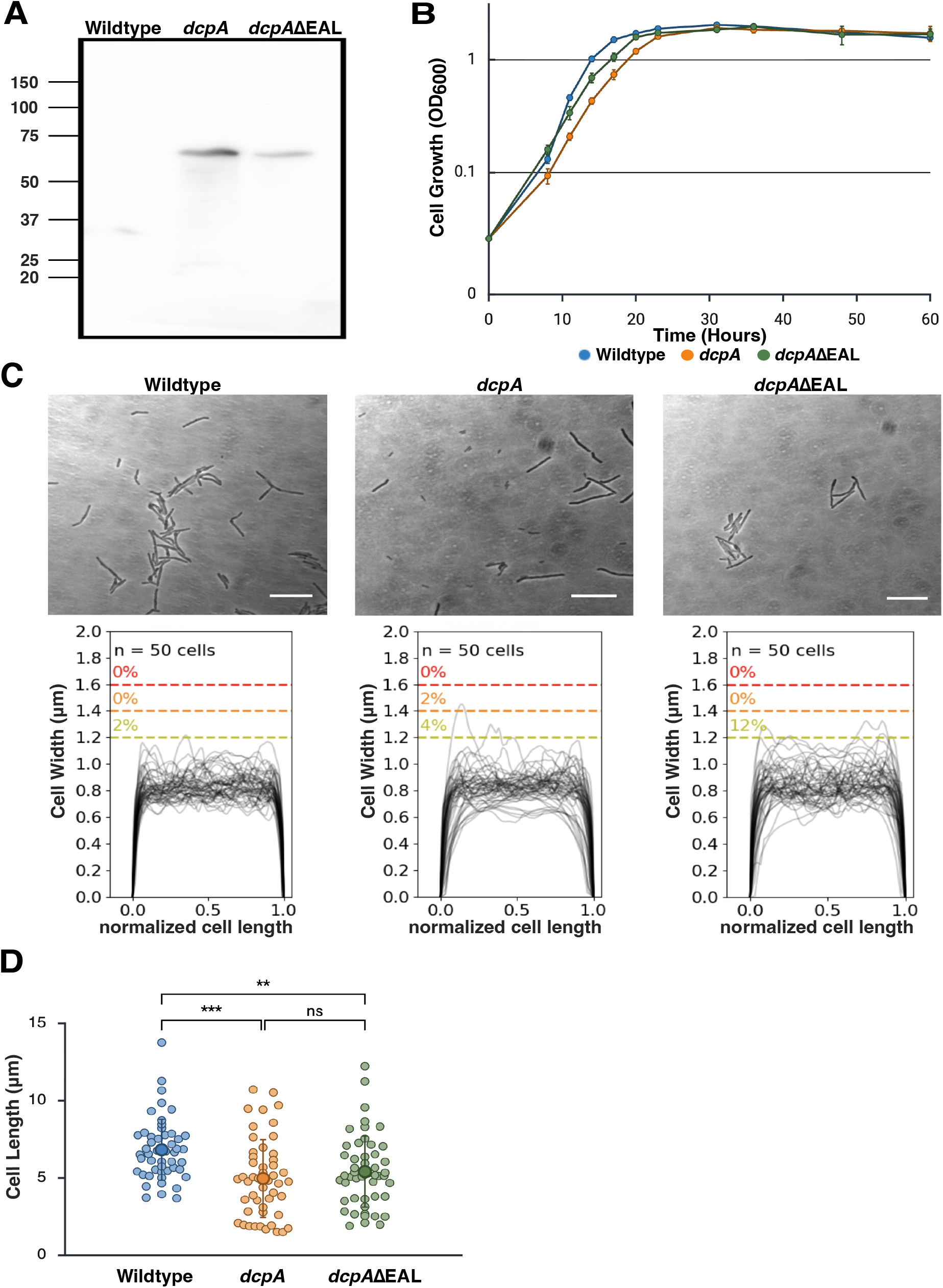
Growth and cell morphology of *dcpA* overexpression *M. smegmatis*. (**A**) Western blot analysis of crude cell lysates prepared from wildtype, *dcpA* overexpression, and *dcpA*ΔEAL overexpression cells. DcpA and DcpAΔEAL were C-terminally tagged with HA epitope, and were detected by an anti-HA antibody. The expected molecular weights of DcpA-HA and DcpAΔEAL-HA are 68.5 and 68.1kDa, respectively. (**B**) Growth curve of wildtype, *dcpA* overexpression, and *dcpA*ΔEAL overexpression strains in Middlebrook 7H9. OD_600_, optical density at 600 nm. (**C**) Phase contrast micrographs and cell width profiles of planktonic wildtype, *dcpA* overexpression, and *dcpA*ΔEAL overexpression strains grown in Middlebrook 7H9. Bars = 5 μm. Cell width profiles were graphed across the normalized cell length for 50 randomly selected cells. (**D**) Dot plot of cell length distributions of wildtype, *dcpA* overexpression, and *dcpA*ΔEAL overexpression strains grown in Middlebrook 7H9. The same 50 cells analyzed in panel C were analyzed for the cell length. Statistical significance was assessed using a Kruskal–Wallis test followed by nonparametric pairwise comparisons using the Steel–Dwass method (α = 0.05). n = 50 cells per group. Wildtype differed significantly from *dcpA* (***, *P* < 0.0001) and *dcpA*ΔEAL (**, *P* = 0.0025), whereas *dcpA* and *dcpA*ΔEAL were not significantly different (ns).

In *Pseudomonas aeruginosa*, elevated c-di-GMP levels redirect cellular resources toward EPS production and lower metabolic activity (Lichtenberg et al., 2022). We therefore tested the possibility that high c-di-GMP levels in our *M. smegmatis* strains could reduce their growth rate. We grew the cells with shaking at 130 rpm in Middlebrook 7H9 containing 0.05% (v/v) Tween-80, a conventional detergent typically added at this concentration to disperse mycobacterial granules. As shown in **Fig. 1B**, strains producing DcpA and DcpAΔEAL have a slight growth defect during exponential growth. However, these overexpression mutants recover from this transient delay and reach comparable cell densities to wildtype during the stationary phase. Next, we examined the cell morphology and found no gross morphological differences among logarithmically growing wildtype, *dcpA*, and *dcpA*ΔEAL overexpression strains (**Fig. 1C**). Cell width profiles were generated across the normalized cell length for 50 randomly selected cells per strain and showed comparable distributions (**Fig. 1C**). A previous study reports that *dcpA* overexpression reduces cell length in *M. smegmatis* (Gupta et al., 2016). Therefore, we next determined the cell lengths of the same 50 cells per group that were analyzed in Fig. 1C (**Fig. 1D**). Nonparametric statistical analysis using a Kruskal–Wallis test followed by Steel–Dwass pairwise comparisons showed significant differences between wildtype and *dcpA* (***, *P* < 0.0001) and between wildtype and *dcpA*ΔEAL (**, *P* = 0.0025), whereas no significant difference was observed between the two overexpression strains.

As c-di-GMP mediates biofilm formation in mycobacteria (Gupta et al., 2015; Gupta et al., 2016; Ling et al., 2024), we tested if these overexpression strains have distinct biofilm phenotypes. We first examined colony size and morphologies of wildtype, *dcpA*, and *dcpA*ΔEAL grown on Middlebrook 7H10 or LB agar. Colonies were grown for 3 days at 37°C and measured for the colony area (**Fig. 2A**). On Middlebrook 7H9 agar, analysis of 42 colonies per strain revealed a subtle but statistically significant difference in colony area between the wildtype and *dcpA* overexpression strains (**, *P* = 0.0273), whereas comparisons involving the *dcpA*ΔEAL strain were not significant (**Fig. 2B**). On LB agar, no significant differences in colony area were observed among any of the strains grown on LB agar, based on measurements of 40 colonies per group (**Fig. 2C**). Statistical significance was assessed using a Kruskal–Wallis test followed by Dunn’s post hoc multiple comparisons test.

**Figure 2.**
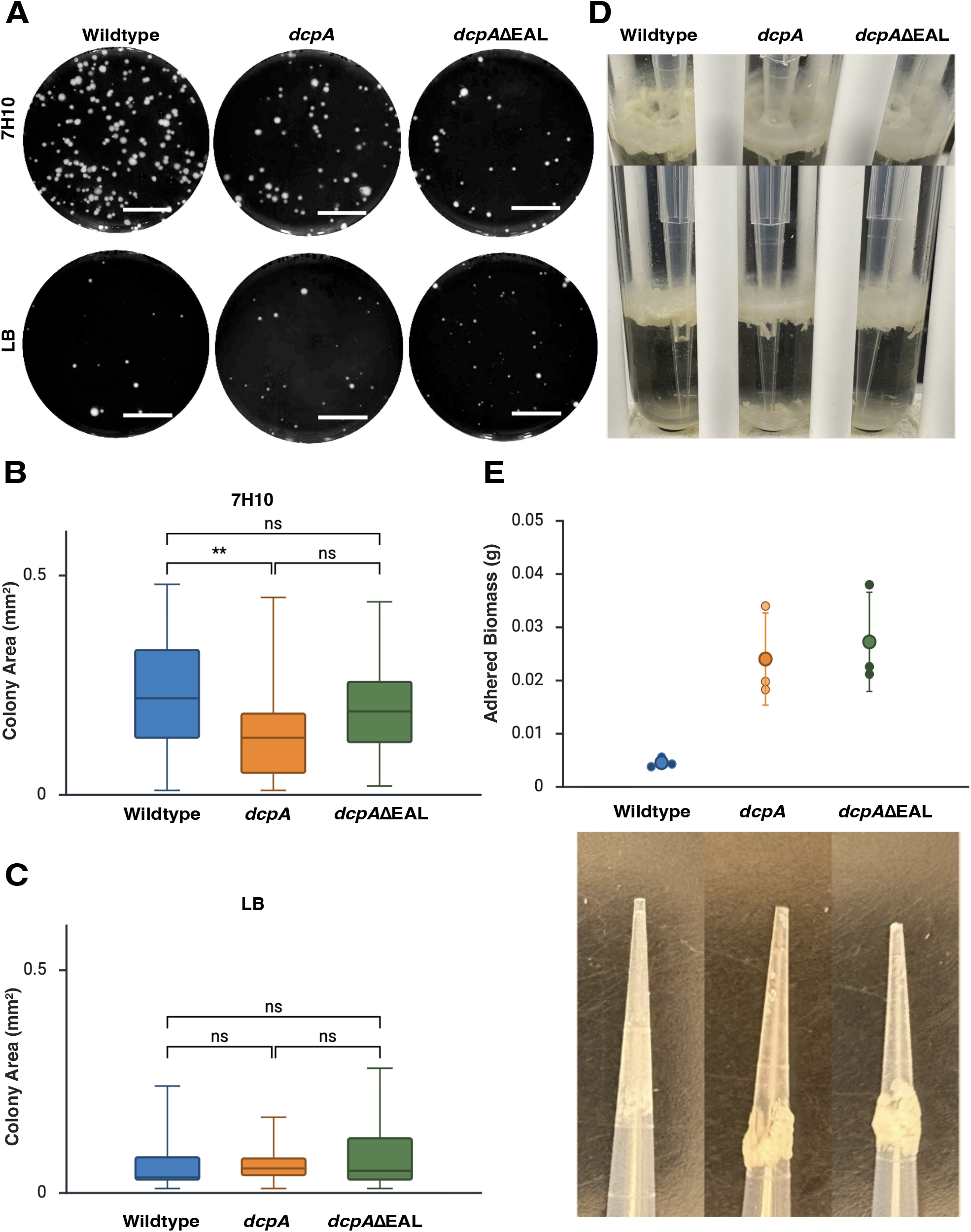
Biofilm properties of *dcpA* overexpression *M. smegmatis*. (**A**) Photographs of colonies of wildtype, *dcpA* overexpression, and *dcpA*ΔEAL overexpression cells grown on Middlebrook 7H10 or LB agar plates. Bars = 5 mm. (**B and C**) Boxplots comparing colony areas among wildtype, *dcpA* overexpression, and *dcpA*ΔEAL overexpression cells grown on Middlebrook 7H10 (B) and LB agar (C). ) A box is drawn from the first quartile to the third quartile with a horizontal line being the median. Whiskers extended to the minimum and maximum observed values. Statistical significance was assessed using a Kruskal–Wallis test followed by Dunn’s post hoc multiple comparisons test. n = 42 or 40 colonies per group for panels B and C, respectively. On 7H10, *dcpA* differed significantly from wildtype (**, *P* = 0.0273), whereas comparison involving *dcpA*ΔEAL were not significant (ns). On LB, comparison among all groups was not significant (ns). (**D**) Top and side views of the surface pellicle with a Pipetman tip co-incubated under static conditions to assess adherence. (**E**) Dot-plot comparing the mass of biofilm adhered on pipette tips from panel B, and representative photo of each tip after removing from glass test tube. Error bars represent the standard deviation of three biological replicates.

### Effects on biofilm phenotype

We then grew overexpression strains in M63 medium (a standard biofilm medium) in the absence of shaking and examined the liquid surface pellicle formation (**Fig. 2D**). Surprisingly, we found that *dcpA* and *dcpA*ΔEAL overexpressing strains showed smooth, glossy surface pellicles without wrinkles, whereas the wildtype formed a robust pellicle with a wrinkled surface, a well-known *M. smegmatis* phenotype. Inspired by a plastic surface adhesion assay reported recently (Ling et al., 2024), we examined whether the mutants, having glossy and sticky appearance, are more adherent to the surface of a polypropylene Pipetman tip (**Fig. 2E**). We observed that the mutants adhered more severely to the plastic tips compared to wildtype, supporting the idea that the DcpA overexpression mutants produce stickier biofilm. Together, our data suggest that these DcpA-expressing strains create a surface architecture that is distinct from that of the wildtype cells.

### Growth and settling behavior in the absence of detergent

Addition of Tween-80 into culture media is an old laboratory practice to alleviate aggregate formation during the growth of *Mycobacterium tuberculosis* and other mycobacteria (Dubos and Davis, 1946; Lyon et al., 1963; Pierce et al., 1947; Power and Hanks, 1965). To test the impact of *dcpA* overexpression on mycobacterial granulation, cells were grown in shaking liquid culture in the absence of Tween-80. When cells are cultured in Middlebrook 7H9 medium without Tween-80, they aggregate and settle, so optical density measurements become unreliable for assessing growth. We therefore analyzed the growth of all three strains by biomass-protein measurement as previously established (Meyers et al., 1998) (**Fig. 3A**). Similar to the growth in Middlebrook 7H9 with the presence of Tween-80, strains overproducing DcpA and DcpAΔEAL showed a slight growth delay during log phase but reached a comparable biomass level to wildtype in stationary phase.

**Figure 3.**
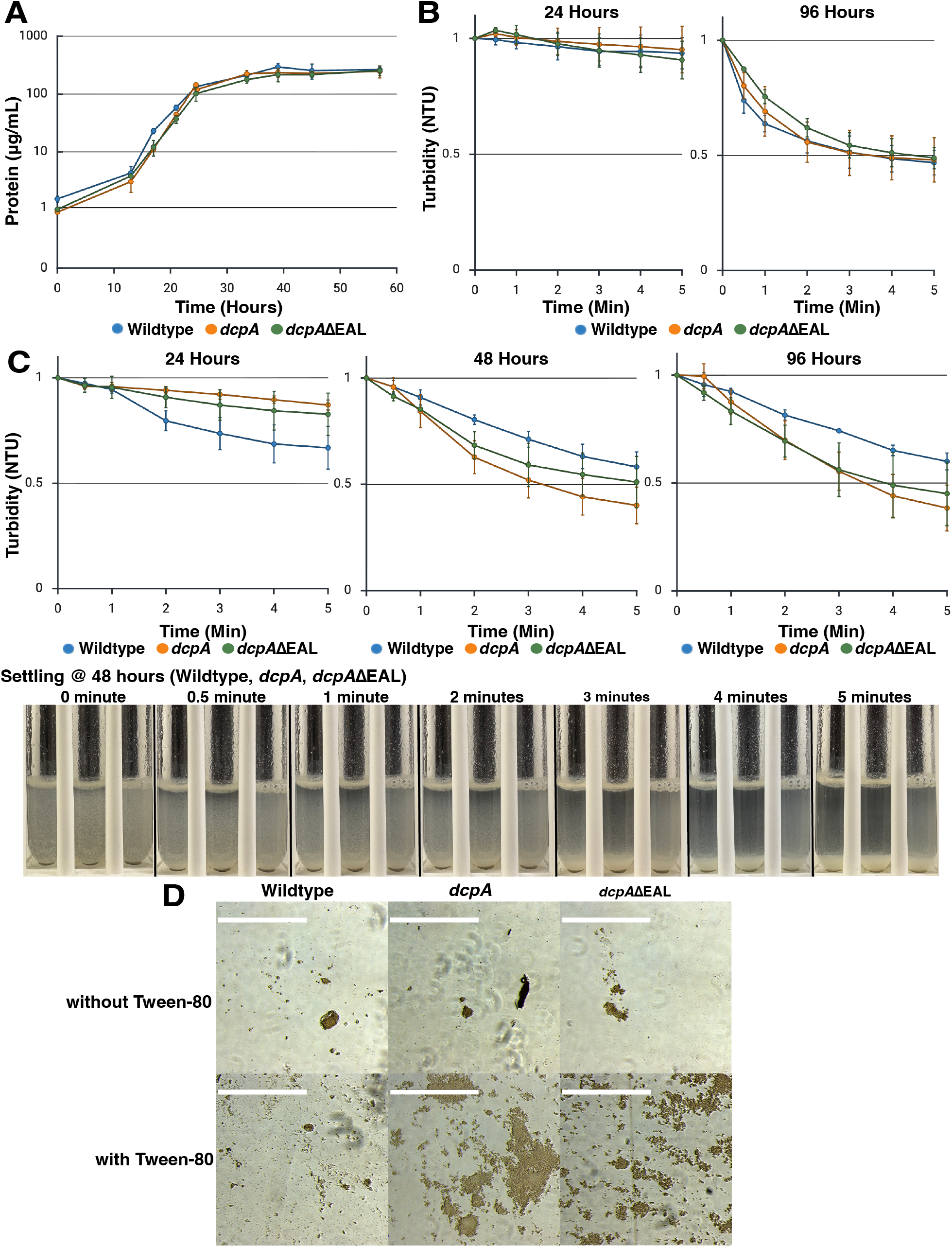
Granular growth and settling properties of wildtype, *dcpA* overexpression, and *dcpA*ΔEAL overexpression *M. smegmatis*. (**A**) Biomass growth curve of wildtype, *dcpA* overexpression, and *dcpA*ΔEAL overexpression cells grown without Tween-80 to promote aggregation in Middlebrook 7H9. Biomass was measured at each timepoint to determine the growth kinetics (see Methods). Error bars represent the standard deviation of three biological replicates. (**B**) Five-min settling of wildtype, *dcpA* overexpression, and *dcpA*ΔEAL overexpression cells grown for 24 and 96 hours in Middlebrook 7H9 with no Tween-80 supplementation. Settling, shown in NTU, was normalized to time 0. Error bars represent the standard deviation of three biological replicates. (**C**) Five-min settling of wildtype, *dcpA* overexpression, and *dcpA*ΔEAL overexpression cells at 24, 48, and 96 hours in Middlebrook 7H9 (supplemented with 0.005% Tween-80). Representative images of settling using cells grown for 48 hours are shown as an example. Settling, shown in NTU, was normalized to time 0, and error bars represent the standard deviation of three biological replicates. (**D**) Micrographs of aggregates of wildtype, *dcpA* overexpression, and *dcpA*ΔEAL overexpression cells grown for 48 hours in Middlebrook 7H9 with and without 0.005 % Tween-80. Bars = 400 µm.

Given that omission of Tween-80 did not impact cell growth, we compared the settling behavior of both wildtype and mutant cultures at late-log and late-stationary phase (24 and 96 hours). We measured supernatant turbidity after allowing cultures to settle for specified intervals (up to 5 min), expecting that better-settling cultures would yield lower turbidity. As shown in **Fig. 3B**, all cultures showed remarkably faster settling in late-stationary phase, indicating stronger cellular aggregation. Unexpectedly, the DcpA-overproducing mutants showed settling speeds that are not significantly different from that of the wildtype (**Fig. 3B**). We reasoned that the intrinsic aggregation of wildtype *M. smegmatis* is so strong that the effect of overexpressing *dcpA* or *dcpA*ΔEAL on granulation would be difficult to detect by settling assays conducted under non-dispersive (*i*.*e*., detergent-free) conditions.

### Settling behavior and phenotype of cells in the presence of detergent

We performed the next set of settling experiments under mildly dispersive conditions to test if DcpA overproduction helps mutant strains to maintain their granulation phenotype. We grew cells in Middlebrook 7H9 containing 0.005% Tween-80, which is 10 times lower than the standard concentration of Tween-80. We then sampled late-log, stationary, and late-stationary phase (24, 48, and 96 hours) cells and examined their settling properties with both turbidity and visual evaluation of the vial images (**Fig. 3C**). At 24-hour time point, wildtype cultures settled faster, possibly because they entered stationary phase slightly earlier than the mutant cultures following marginally faster log-phase growth (**Fig. 1B** and **Fig. 3A**). However, it was clear that at 48-hour and 96-hour time points, both mutant strains overproducing DcpA and DcpAΔEAL showed enhanced settling compared to wildtype, suggesting that *dcpA* overexpression helped to maintain granulation under a mildly dispersive, detergent-present condition. We imaged cell aggregates at 48 hours by phase contrast microscopy, both in the presence and absence of 0.005% Tween-80 (**Fig. 3D)**. In the absence of Tween-80, both wildtype and mutant strains formed aggregates of similar sizes. In contrast, in the presence of Tween-80, wildtype cells only showed small aggregates, whereas the mutant cells produced larger aggregates. Collectively, the results indicate that *dcpA* and *dcpA*ΔEAL overexpression promotes and sustains aggregation under dispersive conditions induced by Tween-80.

### Enhanced aggregation in carbon-rich media

Aggregation of *M. smegmatis* is favored under carbon-rich and nitrogen-limited conditions (DePas et al., 2019). The rich medium TYEM supplemented with 0.6% glucose is particularly effective at promoting and sustaining aggregated growth (DePas et al., 2019). Therefore, we examined the aggregation phenotype of both wildtype and mutants overexpressing *dcpA* and *dcpA*ΔEAL in this carbon-rich medium. Phase-contrast microscopy results showed that both *dcpA* and *dcpA*ΔEAL overexpressing strains grown for 48 hours in this rich media formed larger and denser granules compared with the wildtype (**Fig. 4A**). We further examined the settling properties of cultures cultivated in this rich media (**Fig. 4B**). Compared with the earlier results from cultivation in 7H9 medium + 0.005% Tween-80, both mutant cell types settled much faster than wildtype even from 24-hour growth point in the presence of 0.005% Tween-80. These results suggest that carbon-rich conditions further enhance granulation and settling behavior of *dcpA* and *dcpA*ΔEAL overexpressing *M. smegmatis* strains.

**Figure 4.**
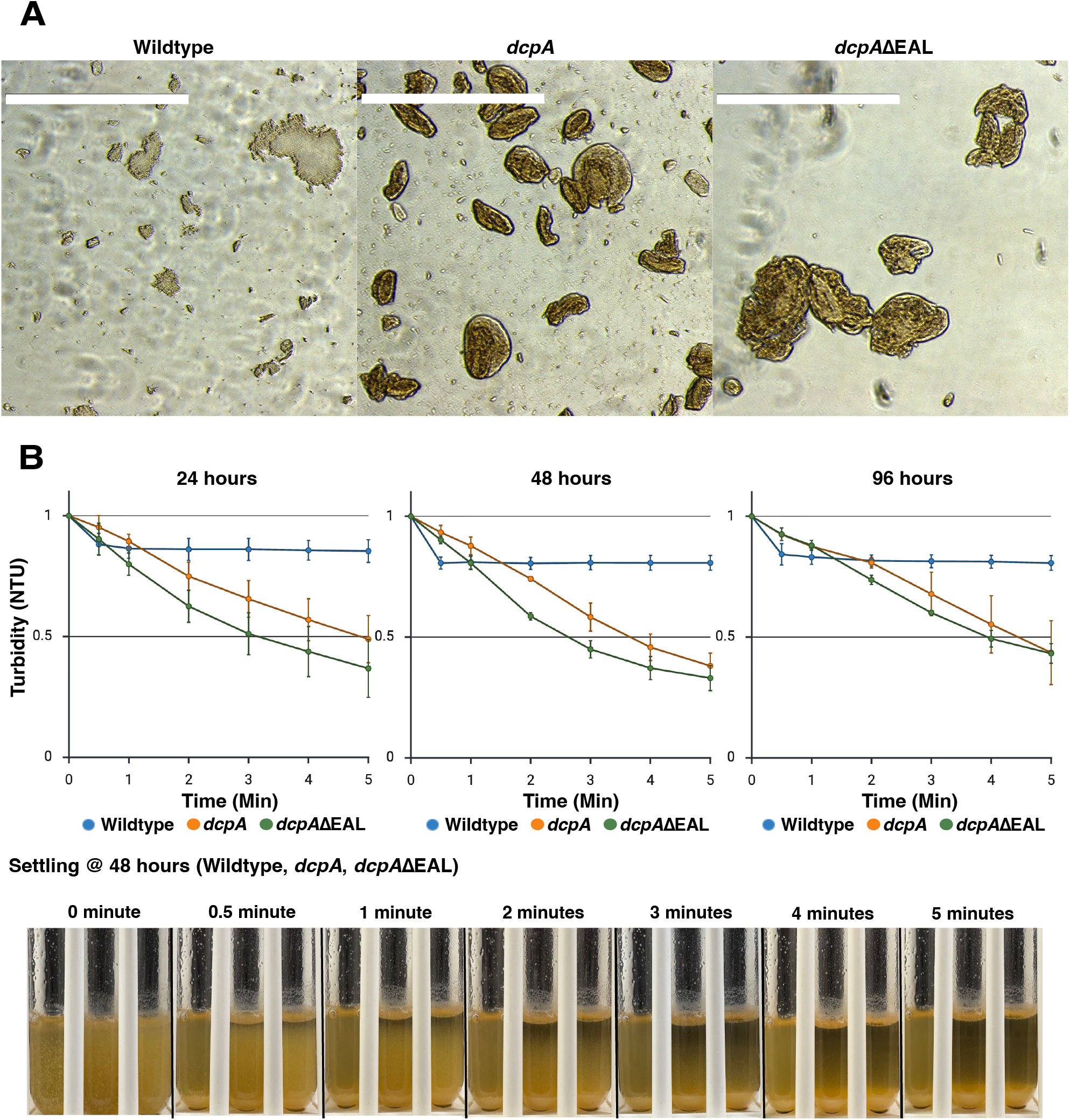
Settling in TYEM medium that promotes cell aggregation. (**A**) Micrographs of aggregates of wildtype, *dcpA* overexpression, and *dcpA*ΔEAL overexpression cells after 48 hours of growth in TYEM medium supplemented with glucose and 0.005 % Tween-80. Bars = 400 µm. (**B**) Five-min settling of wildtype, *dcpA* overexpression, and *dcpA*ΔEAL overexpression cells grown for 24, 48, and 96 hours in TYEM supplemented with glucose (supplemented with 0.005% Tween-80). Representative images of settling using cells grown for 48 hours are shown as an example. Settling, shown in NTU, was normalized to time 0, and error bars represent the standard deviation of three biological replicates.

Granulation typically leads to much lower sludge volume index (SVI) values, such as 30–60 mL/g, than flocculent activated sludge (>100 mL/g), reflecting fundamental differences in biomass structure and settling behavior (Gikonyo et al., 2021; Sarvajith et al., 2024; Wei et al., 2020). To assess whether wildtype and *dcpA*- and *dcpA*ΔEAL-overexpressing *M. smegmatis* form compact granules comparable to granules used in wastewater treatment, we measured their SVI **(Table 1)**. The cells were grown for 48 hours in either Middlebrook 7H9 medium or TYEM medium supplemented with 0.6% glucose. As before, cells were cultivated with 0.005% Tween-80 to apply mild dispersion stress conditions. In both media, both SVI5 and SVI30 were higher for *dcpA*- and *dcpA*ΔEAL-overexpressing strains than the wildtype. This result may appear to contradict the turbidity-based settleability (**Figs. 3C** and **4B**) because SVI reflects only settled biomass volume and does not capture the residual suspended turbidity. Nonetheless, SVI5 and SVI30 for all samples were well below the 100 mL/g threshold, indicating that *M. smegmatis* aggregates fall within the range of compact granules. Moreover, the SVI5/SV30 ratios, which are typically greater than 1 for flocculent sludge (Gikonyo et al., 2021), were all below 1 (**Table 1**), further supporting that *M. smegmatis* forms granules rather than conventional flocs.

**Table 1.** SVI of *M. smegmatis* aggregates. Cells were grown for 48 hours in either Middlebrook 7H9 (7H9) or TYEM supplemented with 0.6% glucose (TYEM) in the presence of 0.005% Tween-80 as a mild granule dispersion stress. Cell aggregates were then harvested for the SVI measurements as described in the Methods section. OE, overexpression. SVI30 values lower than 100 mL/g is empirically suggested to be granulation while those higher than 100 mL/g is considered flocculation (see text for detail).

| Strain | Time (hours) | Medium | SVI5 (mL/g) | SVI30 (mL/g) | SVI5/SVI30 |
| --- | --- | --- | --- | --- | --- |
| Wildtype | 48 | 7H9 | 11 | 19 | 0.677 |
| *dcpA* OE | 48 | 7H9 | 24 | 37 | 0.656 |
| *dcpA*ΔEAL OE | 48 | 7H9 | 19 | 27 | 0.694 |
| Wildtype | 48 | TYEM | 11 | 14 | 0.833 |
| *dcpA* OE | 48 | TYEM | 19 | 24 | 0.810 |
| *dcpA*ΔEAL OE | 48 | TYEM | 20 | 24 | 0.833 |

## Discussion

In this study, we demonstrated that genetic manipulation of DcpA, the native *M. smegmatis* enzyme solely responsible for c-di-GMP synthesis and degradation, alters granulation properties. We showed that overexpression of both *dcpA* and *dcpA*ΔEAL in *M. smegmatis* caused only minimal growth defects (**Figs. 1B and 3A**) and that its effect on granulation was most pronounced when Tween-80 was included to promote dispersion (**Fig. 4**). Based on microscopy and settling assays (**Figs. 3 and 4**), we found that self-aggregates of *M. smegmatis* formed in shaking liquid culture exhibit characteristics with granules rather than flocs. These results indicate that *M. smegmatis* is a tractable model for manipulating c⍰di⍰GMP signaling and bacterial aggregation, offering a platform to develop strategies for optimizing granulation technology.

The mutants overexpressing *dcpA* or *dcpA*ΔEAL exhibited faster settling in 5 min compared to wildtype (**Figs. 3C and 4B)**, suggesting stronger aggregation that the wildtype. Similar enhancements of cellular aggregation and settling were observed in *M. smegmatis* that were genetically modified to express a DGC gene *ydeH* from *E. coli* (Ling et al., 2024). Intriguingly, it was previously reported that a *dcpA* deletion mutant (Δ*dcpA*) of *M. smegmatis* also showed enhanced aggregation in Middlebrook 7H9 supplemented with glucose as a carbon source (Gupta et al., 2015), which may be counteractive to our finding that overexpression of *dcpA* promotes aggregation. Nevertheless, the Δ*dcpA* strain was less adherent to a plastic surface (Gupta et al., 2015) while our *dcpA* overexpression strains as well as *ydeH*-overexpressing *M. smegmatis* showed enhanced plastic surface adhesion compared to wildtype controls (Ling et al., 2024). Despite increased aggregation and adhesion, our strains showed less robust pellicle formation, which is in contrast to *ydeH*-overexpressing *M. smegmatis* that showed a pellicle with denser wrinkles (Ling et al., 2024). These observations may highlight the complex mechanism of bacterial cell aggregation. We currently do not know what causes these phenotypic differences, but speculate that they were attributed to the differences in the properties of the *M. smegmatis* and *E. coli* enzymes. Finally, while *dcpA* overexpression had limited impacts on the growth rate, cell length became shorter. This phenotype is consistent with a previous observation (Gupta et al., 2016).

Given the cellular phenotypes of our *dcpA* overexpression strains, which broadly agree with prior studies on DcpA in *M. smegmatis*, we took a step further to investigate the impact of *dcpA* and *dcpA*ΔEAL overexpression on microbial aggregation with a broader aim of establishing a model for wastewater treatment technologies. Importantly, *dcpA*- and *dcpA*ΔEAL-overexpressing strains sustained more robust aggregation and settleability properties under Tween-80 surfactant stress than the wildtype. Tween-80 is a nonionic detergent routinely added to mycobacterial growth media, typically at 0.05% to prevent cell aggregation. At a concentration of 0.005% Tween-80, the settleability of wildtype strain was markedly reduced compared to both mutant strains. Mounting evidence indicates that c-di-GMP stimulates EPS production in bacteria, thereby providing structural support for biofilm formation (Chen et al., 2025; Cotter and Stibitz, 2007; Isenberg and Mandel, 2024; Purcell and Tamayo, 2016; Römling et al., 2013). EPS can exist as either “loosely bound” or “tightly bound,” each contributing differently to cell survival and the macroscopic appearance of the cell aggregation (Mahto et al., 2022; Wang and Wang, 2023). We postulate that overexpression of *dcpA* and *dcpA*ΔEAL would promote tighter EPS binding, resulting in denser granules that resist Tween-80-induced dispersion and settle effectively. These results also imply that overexpression of *dcpA* and *dcpA*ΔEAL alters cell surface properties, which can be corroborated with a prior study reporting *M. smegmatis* overexpressing *dcpA* produced more EPS than the wildtype as determined by crystal violet staining (Gupta et al., 2016).

In conclusion, our study suggests that intracellular signaling mediated by cyclic-di-GMP promotes stress-resilient granulation in *M. smegmatis*, establishing a useful model of bacterial aggregation in the context of granular technology. By understanding the mechanisms that govern microbial granulation, we can manipulate these traits and improve granulation in wastewater treatment.

## Materials and Methods

### Construction of expression vectors

The *M. smegmatis dcpA* overexpression strain and its ΔEAL variant were made using an integrative expression vector pMUM111 (Puffal et al., 2025), which is based on pMV361 expression vector driven by the strong HSP60 promoter (Stover et al., 1991). The *dcpA* (MSMEG_2196) coding sequence was amplified from *M. smegmatis* genomic DNA using primers carrying NdeI and PacI restriction sites (underlined) for ligation (forward: 5′- AAATAAACCCATATGTCCGAGAGCCTGGACGTGC-3′; reverse: 5′- AAATAAATTTAATTAACATGCGAGGCGCAG CTGAG-3′), digested with NdeI and PacI, and ligated into pMUM111, which was digested with the same enzymes. The resulting construct was transformed into TOP10 chemically competent *E. coli* (Thermo Fisher Scientific), and a correct clone was identified and confirmed by restriction enzyme digestions and Sanger sequencing, resulting in the *dcpA* expression vector designated as pMUM409. To create an in-frame deletion of the EAL motif, we used the Q5 Site-Directed Mutagenesis Kit (New England Biolabs). We designed primers 5′-GTCCGCTGGGAGCATCCCACC-3′ and 5′-TGCCGCCAGCACCTCACC-3′, and deleted the EAL motif using pMUM409 as the template. The deletion was verified by Sanger sequencing, resulting in pMUM410. The plasmids were electroporated into wildtype *M. smegmatis* mc^2^155 using standard protocols (Morita et al., 2006) except that ECM 830 Square Wave Electroporation System (BTX) was used, and kanamycin-resistant transformants were isolated.

### Growth

For planktonic growth of *M. smegmatis* mc^2^155, cells were grown in Middlebrook 7H9 supplemented 11□mM glucose, 14.5□mM NaCl, and 0.05% (v/v) Tween-80. Kanamycin was added at 20 µg/ml for growing seed cultures of overexpression strains to ensure the maintenance of the expression vector. To facilitate cell aggregation, cells were grown without Tween-80 addition. To challenge cells with aggregation dispersion stress, cells were grown with 0.005% (v/v) Tween-80. For enhanced aggregation, cells were grown in TYEM (0.5% (w/v) tryptone, 0.25% (w/v) yeast extract, and 2 mM MgSO_4_) supplemented with 0.6% glucose and 0.005% (v/v) Tween-80 (DePas et al., 2019). For pellicle biofilm growth, 30 µL planktonic stationary phase culture was diluted 1:100 in 3 mL of M63 medium in glass test tubes and incubated for 5 days at 37°C without shaking. Images of pellicles and biofilm morphology were recorded using an iPhone 16 camera (Apple). For colony growth, serially diluted planktonic cultures were spread evenly on Middlebrook 7H10 and LB solid agar plates, incubated for 3 days at 37°C. Colonies were imaged using Amersham ImageQuant 800 (Cytiva) and colony area was determined using ImageJ software (Schneider et al., 2012). Statistical analysis of colony area was performed using JMP Student Edition (version 18).

### Biomass measurement

To assess growth by biomass measurement, 100 mL of Middlebrook 7H9 without 0.05% Tween-80 in a 500-mL glass flask was inoculated with seed culture to a starting OD_600_ of 0.02. At each time point, roughly 2 OD units was collected, spun down at 3,220x *g* for 10 mins (Eppendorf 5810R) or 16,900x *g* for 2 mins (Eppendorf 5418R). Supernatants were removed and pellets were washed in 1 mL of PBS. Pellets were stored at 20°C until all samples were collected. For protein extraction, 50 µL of NaOH was added to pellets, sonicated to disperse the cell pellets, and incubated for 10 min at 95°C. To neutralize pellets, 8.33 µL of 6 M HCl was added. Volume was adjusted to 0.5 mL by adding PBS and then transferred to 1.5 mL microtube. Pellets were spun down for 30 min at 16,900x g (Eppendorf 5418R). Using 1 µL of supernatant, absorbance was read at 230 and 260 nm using Nanodrop One C UV-Vis Spectrophotometer (ThermoFisher). Protein concentration (μg/ml) was calculated as follows: [Protein] = (183 × A_230_) - (75.8 × A_260_)

### SDS-PAGE and western blotting

Cell lysates were prepared by bead beating as previously described (Rahlwes et al., 2017). Protein samples were mixed with reducing loading dye and denatured by boiling for 5 min at 95°C. Samples were separated on SDS-PAGE (10% separating gel) at 110 V. For western blotting, gel was transferred onto polyvinylidene difluoride (PVDF) membrane, sandwiched between Whatman papers (Bio-Rad) in a transfer buffer (25□mM Tris-HCl (pH 8.3), 192□mM glycine, 0.1% SDS (w/v)) at 14□V on ice. After overnight transfer, the membrane was blocked for an hour in blocking buffer (5% skim milk in PBS□+□0.05% Tween-20 (PBST)). The membrane was then incubated with the primary mouse anti-HA antibody (Millipore Sigma) used at a final concentration of 0.5 µg/mL for two hours, washed in PBST for 10 min three times, and further incubated with the secondary sheep anti-mouse IgG antibody conjugated with horseradish peroxidase (GE Healthcare) at a 1:2,000 dilution for one hour. The membrane was washed again and developed with homemade chemiluminescence reagent (2.24□mM luminol, 0.43□mM *p*- coumaric acid, and 0.0036% H_2_O_2_ in 100□mM Tris-HCl (pH 9.35)). The chemiluminescence signals were recorded using Amersham ImageQuant 800 (Cytiva).

### Microscopy and morphological analysis

Log and stationary phase cells were spotted onto agar pads (1% agarose gel) placed on a glass slide and imaged at 1000x magnification with Nikon Eclipse E600 microscope (100x objective; numerical aperture, 1.30) equipped with an ORCA-ER cooled charge-coupled-device camera (Hamamatsu). Cell morphology was analyzed and graphed as before using Oufti (Paintdakhi et al., 2016) and a previously published custom python script (Sparks et al., 2024). For microscopy of aggregates, cells were grown for 48 hours in Middlebrook 7H9 or TYEM supplemented with 0.6% glucose with and without 0.005% Tween-80. The culture flasks were settled for 5 min before 10 μl of sample was harvested from the bottom of the flask and placed onto a glass slide. The aggregates were viewed and imaged at 10X magnification using a bright field light microscopy (EVOS FL Color, ThermoFisher, AMEFC-4300) to characterize morphology. All experiments were performed in biological triplicate.

### Adherence assay

To assess adhesion properties, 20 mL of planktonic seed cultures of each strain were grown as described above with shaking for 3 days at 37°C. For adhesion assays, 3 mL of sterile M63 medium was aliquoted into glass test tubes and inoculated with 30 µL of the corresponding seed culture. A sterile 200 µL Pipetman tip, pre-weighed prior to incubation, was vertically submerged into each tube. Cultures were incubated statically at 37°C for 5 days. Following incubation, pellicle formation was visually assessed and documented. Tips were carefully removed, imaged to document attached biomass, and re-weighed to quantify bacterial adhesion by mass difference. Experiment was performed in biological triplicate.

### Settling property assay

To test settling properties, 10 mL of *M. smegmatis* cells grown in Middlebrook 7H9 or TYEM supplemented with 0.6% glucose with or without 0.005% Tween-80 at 24, 48, and 96 hours was aliquoted, and placed into 1” Round Glass vials (10 mL volume, Hach). Vials were inverted twice, immediately placed into DR900 Multiparameter Portable Colorimeter (Hach), and measurement was immediately taken for time 0. Measurement was subsequently taken at 0.5, 1, 2, 3, 4, and 5 min. Settling, as measured by nephelometric turbidity unit (NTU), was normalized to show settling relative to the 0-min time point. All experiments were performed in biological triplicate.

### SVI assay

Cells were grown with 0.005% Tween-80 as described above. After 48 hours of shaking at 37 °C, 10 mL of culture was transferred to a graduated cylinder, mixed by inverting three times, and total volume (Vs) recorded. Settled volumes were measured at 5 min (SV5) and 30 min (SV30). Because TYEM medium is a rich medium that skews SVI values, we modified the SVI calculation to use settled solids (SS) instead of total solids (TS). To measure SS, empty aluminum trays were baked at 105 °C for 30 min, cooled in a desiccator for 30 min, and weighed to record the tray weight (M_1_). Ten mL of cultures were then centrifuged at 3,220x *g* for 10 min, washed with 1 mL water, and the pellet resuspended in 500 µL water. The suspended cells were then transferred to the tray and dried at 105°C for up to 1.5 hours until the weight becomes constant. The final weight of the tray with the dried biomass was recorded (M_2_). Experiment was performed in triplicate, and SS, SVI5, and SVI30 were calculated as follows: SVI = Settled Volume (mL) / SS, where SS (g/L) = (M_2_ – M_1_) / Vs.

## Acknowledgements

We thank Laney Celli and Paige Bosworth for technical help. This study was partly supported by the Engineering Faculty Research Initiative Fund from the Riccio College of Engineering at the University of Massachusetts Amherst (to C.P.).

